# Pan-genome study underlining the extent of genomic variation of invasive *Streptococcus pneumoniae* in Malawi

**DOI:** 10.1101/2023.01.02.522535

**Authors:** Arash Iranzadeh, Arghavan Alisoltani, Anmol M Kiran, Robert F Breiman, Chrispin Chaguza, Chikondi Peno, Jennifer E Cornick, Dean B Everett, Nicola Mulder

## Abstract

**Abstract:** *Streptococcus pneumoniae* is a common cause of acute bacterial infections in Malawi. Understanding the molecular mechanisms underlying its invasive behavior is crucial for designing new therapeutic strategies. We conducted a pan-genome analysis to identify potential virulence genes in *S. pneumoniae* by comparing the gene pool of isolates from carriers’ nasopharyngeal secretions to isolates from the blood and cerebrospinal fluid of patients. Our analysis involved 1,477 pneumococcal isolates from Malawi, comprising 825 samples from carriers (nasopharyngeal swab) and 652 from patients (368 from blood and 284 from cerebrospinal fluid). We identified 56 serotypes in the cohort. While most serotypes exhibited a similar prevalence in both carriage and disease groups, serotypes 1 and 5, the most abundant serotypes in the entire cohort, were significantly more commonly detected in specimens from patients compared to the carriage group. This difference is presumably due to their shorter nasopharyngeal colonization period. Furthermore, these serotypes displayed genetic distinctiveness from other serotypes. A magnificent genetic difference was observed in the absence of genes from the RD8a genomic island in serotypes 1 and 5 compared to significantly prevalent serotypes in the nasopharynx. RD8a genes play pivotal roles in binding to epithelial cells and performing aerobic respiration to synthesize ATP through oxidative phosphorylation. The absence of RD8a from serotypes 1 and 5 may be associated with a shorter duration in the nasopharynx, theoretically due to a reduced capacity to bind to epithelial cells and access free oxygen molecules required for aerobic respiration (essential to maintain the carriage state). Serotypes 1 and 5, significantly harbor operons that encode phosphoenolpyruvate phosphotransferase systems, which might relate to transporting carbohydrates, relying on phosphoenolpyruvate as the energy source instead of ATP. In conclusion, serotypes 1 and 5 as the most prevalent invasive pneumococcal strains in Malawi, displayed considerable genetic divergence from other strains, which may offer insights into their invasiveness and potential avenues for further research.

**Author summary:** Despite introducing the pneumococcal conjugate vaccine in 2011, *Streptococcus pneumoniae* remains a major cause of bacterial infection in Malawi. Whilst some pneumococcal strains harmlessly colonize the nasopharynx, others find their way into normally sterile sites, such as lungs, blood, and nervous system, resulting in serious illness. Our study identified specific pneumococcal serotypes as the most invasive in Malawi, characterized by a short colonization period and significant genetic distinctiveness from other strains. This genetic divergence notably included the absence of several genes associated with aerobic respiration and the presence of genes facilitating ATP-independent carbohydrate transport. The presence or absence of these genes may underlie their heightened invasiveness and shorter colonization period. This hypothesis positions these genes as potential candidates for future therapeutic research. We propose that the specific gene gain and/or loss in invasive versus other serotypes may be linked to the development of invasive pneumococcal diseases.

**Impact Statement:** Our research applied pan-genomics principles to comprehensively assess diversity within the pneumococcus genome, with the primary objective of identifying pneumococcal virulence genes for advancing vaccine design and drug development. Within this study, we identified Serotypes 1 and 5 as the predominant and highly invasive pneumococcal strains in Malawi, characterized by a short nasopharyngeal colonization period, suggesting their potential for rapid infection of sterile sites within the human body such as blood and the central nervous system. These serotypes exhibited significant genetic divergence from other serotypes in Malawi, notably lacking key genes within the RD8a operon while harboring transporters functioning independently of ATP. It’s important to note that these findings are based on computational analysis, and further validation through laboratory experiments is essential to confirm their biological significance and potential clinical applications. The implications of our research offer potential avenues for more effective pneumococcal disease prevention and treatment, not only in Malawi but also in regions facing similar challenges.

## Introduction

*Streptococcus pneumoniae,* also known as *pneumococcus*, is a Gram-positive, facultatively anaerobic bacteria and is one of the leading causes of mortality worldwide. Despite reductions in the incidence of pneumococcal disease in countries that introduced pneumococcal conjugate vaccines (PCV), the pneumococcal mortality rate is still high. Pneumococci are estimated to be responsible for 317,300 deaths in children aged 1 to 59 months worldwide in 2015 [1]. In the post-PCV era, a high disease burden has still been reported in low-income countries in Africa, such as Malawi [2].

Although pneumococcal nasopharyngeal colonization is asymptomatic, it is a prerequisite for transmission and disease development [3][4]. Symptoms appear when isolates from the nasopharynx spread to normally sterile sites such as the lung, blood, and central nervous system. Depending on the infected organ, *S. pneumoniae* can cause two types of infection: (i) non-invasive (mucosal) pneumococcal diseases such as otitis media and sinusitis, and (ii) invasive pneumococcal diseases (IPD) such as bacteremia and meningitis. IPD incidence is highest among infants, the elderly, and immunosuppressed people, most likely due to their less efficient immune systems [5].

Pneumococci possess several virulence factors, including the polysaccharide capsule, surface proteins, and enzymes [6][7]. The polysaccharide capsule is the most important virulence factor as it aids the *pneumococcus* in evading the immune response during colonization and invasion [8]. Its biosynthesis is regulated by a cluster of genes in the capsular polysaccharide (*cps*) locus [8][9]. Pneumococcal serotypes are defined by the type and order of monosaccharides that compose the capsule structure. To date, one hundred pneumococcal serotypes have been identified [10]. Each strain has a set of capsular synthesis genes in the *cps* locus that determine its serotype. Immunogenic properties of the capsular polysaccharide were utilized to develop all pneumococcal vaccines in use, including PCV7, PCV10, and PCV13 that cover 7, 10, and 13 serotypes, respectively. PCV13 includes the following serotypes: 1, 3, 4, 5, 6A, 6B, 7F, 9V, 14, 18C, 19A, 19F, and 23F. Although the introduction of PCVs has significantly reduced the burden of disease caused by vaccine types (VTs), serotype replacement has increased the non-vaccine types (NVTs) carriage rate and IPD incidence [11][12].

In November 2011, PCV13 was introduced in Malawi, which markedly decreased the health system burden and rates of severe childhood pneumonia [13]. A case-control study in Malawi showed vaccine effectiveness against VT-IPD of 80·7% [14]. A nasopharyngeal carriage survey conducted in the Karonga district showed that although the vaccine reduced the VT colonization rate, a moderate level of serotype replacement was observed among carriers [15]. Moreover, the emergence of antibiotic-resistant pneumococci due to the overuse of antibiotics is a global concern in the 21^st^ century [16]. To develop new, more effective vaccines against the vaccine-escape clones and design effective drugs against antibiotic-resistant strains, it is critical to understand the functions of genes involved in pneumococcal colonization and pathogenesis. During the past decade, the evolution of high-throughput sequencing technologies has generated enormous amounts of genomic data that have enabled researchers to perform large-scale genomic analysis. A well-known example is pan-genome studies. The pan-genome is the entire gene set in a collection of closely related strains within a specie [17]. Determining the genetic drivers is an active and promising area of research that can provide insights into pneumococcal disease prevention and treatment to reduce mortality rates. The pan-genome is useful for analyzing recombinogenic pathogens such as *pneumococcus* [18]. The recombination level is high in thirteen pneumococcal genomic loci known as regions of diversity (RDs) numbered from RD1 to RD13, some of which are involved in virulence [19][20].

In this study, we conducted whole-genome sequencing (WGS) on 1477 pneumococcal samples from residents of Blantyre, Karonga, and Lilongwe in Malawi. Our study aims to: (i) identify serotype distribution, (ii) characterize the pneumococcal population structure, and (iii) identify potential driver genes for invasion and their biological functions.

## Materials and methods

### Study design and sample collection

The study utilized archived samples maintained by the Malawi-Liverpool Wellcome Trust Clinical Research Programme (MLW). Samples were collected from individuals residing in three distinct regions of Malawi: Blantyre in the south, Karonga in the north, and Lilongwe in the central part of the country. This cohort included isolates obtained from both asymptomatic carriers and symptomatic patients.

Carriage samples were collected from the nasopharynx of healthy individuals as part of the Health and Demographic Surveillance System in Karonga and Blantyre. The collection process involved the use of nasopharyngeal swabs. Subsequently, pneumococcal isolates were identified utilizing a previously established protocol [21]. Briefly, the identification method involved culturing isolates on blood agar supplemented with gentamicin, with further confirmation relying on optochin disc-based assays, scrutinizing colony morphology, alpha-hemolysis, and optochin susceptibility, adhering to established norms and practices for pneumococcal isolates. To account for the true diversity of carriages, only a single isolated colony was sequenced and serotyped, therefore no carriage samples included in this study had multiple serotypes.

Invasive pneumococcal samples were also sourced from archived bacterial isolates at MLW, which had been collected from the blood and cerebrospinal fluid (CSF) of symptomatic patients attending Queen Elizabeth Central Hospital in Blantyre and Kamuzu Central Hospital in Lilongwe. Notably, the selection of isolates for this group was blind to their serotypes, ensuring an accurate representation of their prevalence in the disease group without any influence from serotype inclusion criteria. These isolates were subsequently streaked onto blood agar plates supplemented with gentamicin, and optochin tests were conducted, according to the procedures outlined in reference [22].

It’s important to note that this study did not involve paired samples. During data collection, each individual contributed only one sample, which was either a nasopharyngeal swab from healthy individuals or a blood or CSF sample from symptomatic patients. In the context of this study, the term ‘sterile sites’ refers to blood and CSF. The term ‘invasive samples’ specifically refers to those samples obtained from these sterile sites (blood and CSF).

### Whole-genome sequencing and quality control

Archived samples were sequenced under the Global Pneumococcal Sequencing project and Pneumococcal African Genomic Consortium at the Wellcome Trust Sanger Institute in the United Kingdom. Bacterial DNA was extracted using the QIAamp DNA mini kit and QIAgen Biorobot by QIAGEN. Whole-genome sequencing was conducted on Illumina Genome Analyzer II and HiSeq platforms, producing 125 nucleotide paired-end reads. Read quality was assessed using Fastqc [23].

### In-silico serotyping, sequence typing, and quantification of serotype invasiveness

SeroBA version 1.23.4 was employed to infer the serotype of the samples [24]. SeroBA applies a k-mer method to determine serotypes directly from the paired-end reads in FASTQ format. Any serotype with a relative frequency greater than 5% was categorized as an abundant serotype. To identify serotypes with a significant presence in the nasopharynx and sterile sites, Fisher’s exact test was applied. P-values were adjusted using the Benjamini-Hochberg method, and serotypes with an adjusted p-value less than 0.01 were considered significant. The odds ratio (OR) was calculated as follows: OR = (ad)/(bc), where ‘a’ represents the number of invasive serotype k, ‘b’ is the number of carriage serotype k, ‘c’ is the number of invasive non-serotype k, and ‘d’ is the number of carriage non-serotype k. Zero values were replaced by 0.5 in OR calculations. Abundant serotypes with a significant presence in the nasopharynx were considered to have low invasiveness, while abundant serotypes with a significant presence in sterile sites were considered to have high invasiveness. Fisher’s exact test was also used to identify serotypes whose frequencies changed significantly after the introduction of PCV13 in 2011 (adjusted p-value < 0.01).

### Genome assembly and annotation

Genomes were assembled using Velvet Optimiser version 2.2.5 [25] with settings to generate contigs longer than 500 base pairs, employing a hash range from 61 to 119. The quality of the assembled genomes was assessed using Quast version 5.2 [26], and annotation was performed with Prokka version 13.1 [27].

### Pan-genome construction

The pan-genome for the samples was generated using Roary version 3.12.0 [28]. Roary was run to perform the core gene alignments with Mafft version 7.313 [29]. Genes in the pan-genome were categorized into three groups based on their abundance among samples: genes present in 100% of samples were designated as core genes, those in more than 95% but not core were termed soft-core genes, and the remaining genes were considered accessory genes.

### Analysis of the population structure

Small-scale variations, such as single nucleotide polymorphisms (SNPs) and short indels, within the core genes were analyzed to understand population diversity. A phylogenetic tree, illustrating the genetic separation between samples, was constructed using SNPs and indels in the core gene alignment as phylogeny markers. The core genome alignment served as input for IQ-TREE version 2 [30] to generate a phylogenetic tree, which was visualized using iTol version 3 [31].

Diversity in the accessory genome manifests as large-scale gene presence-absence variations. The R package Nonnegative Matrix Factorization [32] was used to create a gene presence-absence heatmap from the pan-genome matrix. To determine the factors influencing gene distribution, including isolation sites (nasopharynx, blood, and CSF), serotypes, geographical locations (Blantyre, Karonga, and Lilongwe), and vaccination eras, a principal component analysis (PCA) of gene distribution was conducted using the R package MixOmics version 6.20.0 [33].

### Gene presence-absence statistical analysis

Phenotypic traits of samples were assigned based on population structure and invasiveness. To identify putative virulence factors, a gene presence-absence statistical test was conducted using Scoary version 1.6.1643 [34]. This tool scores the components of the pan-genome for associations with observed phenotypic traits while accounting for population stratification. The test was conducted across samples from different sources and serotypes to find putative virulence factors. Genes with a Bonferroni-corrected p-value less than 0.05 were deemed significant.

### Functional and gene ontology (GO) enrichment analysis

The list of significant genes was submitted to STRING webtool version 11.5 [35] for functional enrichment analysis. STRING is a network that integrates information from various protein-protein interaction databases, predicting both direct (physical) and indirect (functional) interactions between proteins from five sources, including genomic context predictions, lab experiments, co-expression, automated text mining, and previous knowledge in databases. Functional enrichment analysis in STRING utilizes information from classification systems such as the Kyoto Encyclopedia of Genes and Genomes (KEGG) [36] and the Protein families database (Pfam) [37]. The tool entitled “Multiple Sequences” was selected, and the *S. pneumoniae TIGR4* was chosen as the reference organism. STRING reports the associated enriched pathways with a false discovery rate of less than 0.05.

## Results

In total, 825 isolates from the nasopharynx of healthy carriers, 368 isolates from the blood of bacteremia patients, and 284 isolates from the CSF of meningitis patients were sequenced. The demographics of the samples are shown in Table 1.

**Table 1.** Demographics of 1477 pneumococcal isolates collected from Malawi.

| Characteristics | Categories | Nasopharynx | Blood | CSF |
| --- | --- | --- | --- | --- |
| Age<br>(in years) | < 5 | 538 | 165 | 141 |
|  | 5-19 | 109 | 42 | 60 |
|  | 20-40 | 60 | 67 | 50 |
|  | > 40 | 7 | 24 | 7 |
|  | Unknown | 111 | 70 | 26 |
| Sex | Female | 401 | 131 | 111 |
|  | Male | 313 | 122 | 122 |
|  | Unknown | 111 | 115 | 51 |
| City | Blantyre | 169 | 357 | 259 |
|  | Karonga | 656 | 0 | 0 |
|  | Lilongwe | 0 | 0 | 23 |
|  | Unknown | 0 | 11 | 2 |
| Sampling period |  | 2009-2014 | 1997-2015 | 2000-2015 |

### Serotypes 1, 5, and 12F had the highest invasiveness, likely with a short period of nasopharyngeal colonization

Altogether, we identified isolates belonging to 56 different serotypes. Irrespective of their isolation sources, serotypes 1 (8.7%), 5 (7.8%), 6B (6.6%), 23F (6.3%), and 19F (5.5%) were the most prevalent, each accounting for over 5% of the entire cohort. Of the samples, 66% were collected prior to the introduction of PCV13 in 2011, while 27% were obtained in the post-PCV13 era (S1 Fig). In the pre-PCV13 era, prominent serotypes with frequencies exceeding 5% included 5 (10.21%), 6B (8.72%), 1 (8.4%), 23F (7.23%), 6A (5.74%), and 16F (5.53%). In the post-PCV13 era, serotypes 1 (11.5%) and 12F (5.4%) predominated. It is worth noting that serotype 1 exhibited sustained dominance, with its frequency increasing following the vaccine rollout, while serotype 12F emerged as an abundant strain after 2011 (S2 Fig). Nevertheless, a more extensive dataset encompassing vaccination information could offer further insights into this phenomenon.

Within the carriage isolates, serotypes exhibited abundant frequencies, included 19F (7.88%), 6B (7.27%), 16F (6.79%), and 23F (5.21%), with each surpassing a 5% frequency threshold. The distribution of serotypes among carriers in Blantyre and Karonga largely mirrored each other, with the exception of serotype 13, which displayed higher prevalence in Blantyre, and serotype 6B, which exhibited greater dominance in Karonga (S3 Fig).

Among the blood samples, serotype 5 (20.38%) was the most dominant, followed by 1 (16.58%), 23F (8.42%), and 6B (5.98%). In the cerebrospinal fluid (CSF) samples, serotypes 1 (21.13%), 5 (8.1%), 12F (7.04%), 23F (6.69%), 6A (5.63%), and 6B (5.28%) predominated. When considering the combined blood and CSF groups, the most frequently observed serotypes were 1 (18.56%), 5 (15.03%), 23F (7.67%), 6B (5.67%), and 6A (5%). It’s noteworthy that the majority of invasive samples were collected in Blantyre (96.5%). The invasive samples from Lilongwe were exclusively CSF samples, with serotypes 1 and 12F being dominant (S4 Fig).

As depicted in Fig 1 and detailed in Supplementary Table 1 (S1 Table), among the serotypes that were abundant in either the blood or CSF, serotypes 1 (p = 1.96E-34), 5 (p = 3.97E-19), and 12F (p = 5.29E-06) exhibited a significant presence among patients, with a low frequency of occurrence in the nasopharynx. Given that the colonization phase is a prerequisite for infection, it is plausible that serotypes 1, 5, and 12F may have a short period of nasopharyngeal colonization before infecting sterile sites. Considering that serotypes 1 and 5 were also the most prevalent across the entire cohort, they could be regarded as the most common serotypes with the highest invasiveness. In this study, we categorize serotypes 1, 5, and 12F as ‘significant invasive serotypes’ or ‘hyper-invasive serotypes’.

**Fig 1.**
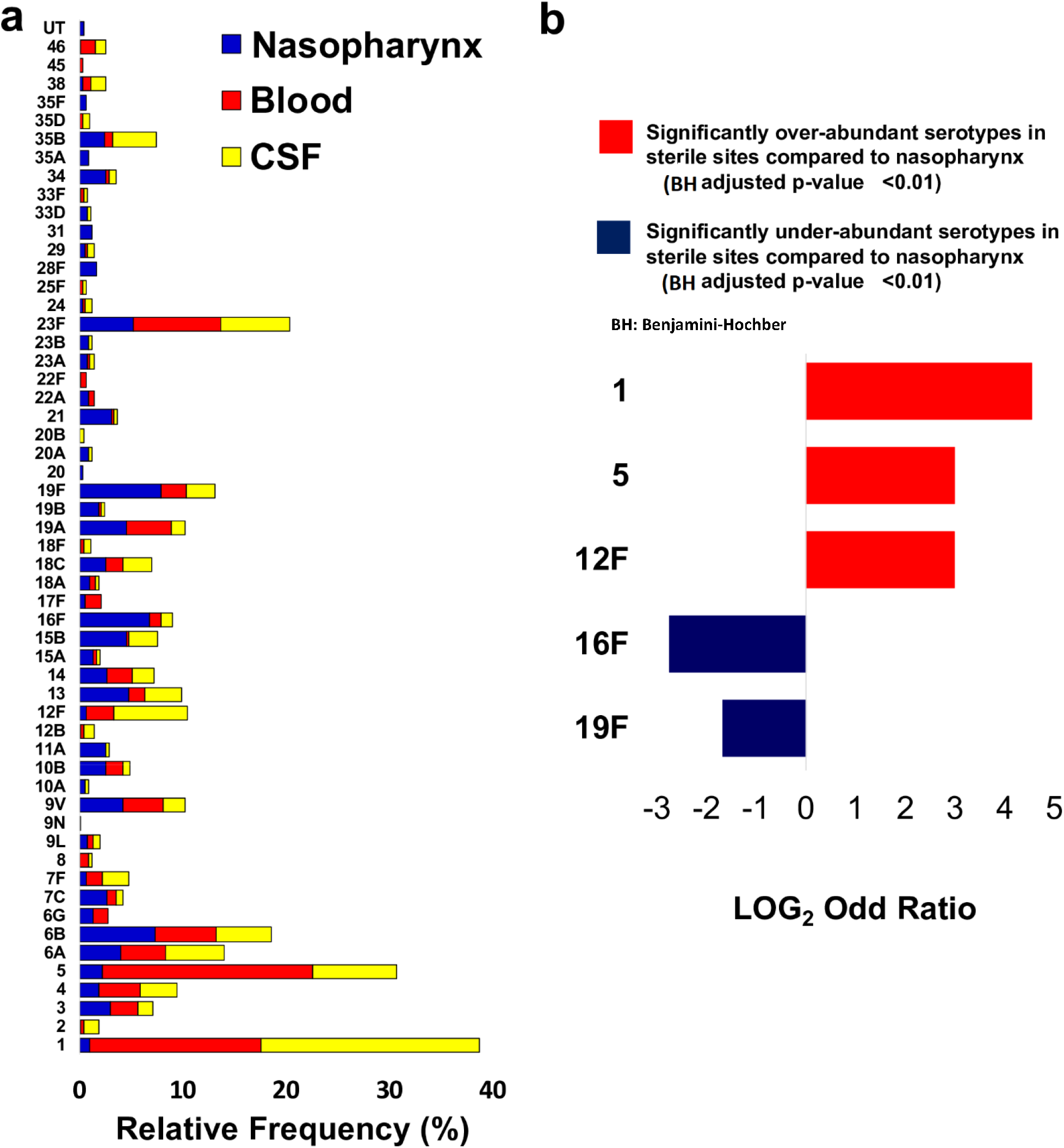
The distribution of the 56 pneumococcal serotypes assigned to 1477 samples from Malawi. (a) The relative frequency of each serotype in the nasopharynx of carriers, the blood of bacteremia patients, and the CSF of meningitis patients is shown in blue, red, and yellow, respectively (UT: Un-Typeable). (b) The log-transformed odds ratio of the significantly over- and under-abundant serotypes in the sterile sites (blood and CSF). Fisher’s exact test was applied to identify serotypes with a significant differential abundance among carriers and patients (nasopharynx and sterile sites) at the significance level of the Benjamini-Hochberg adjusted p-value < 0.01 (BH: Benjamini-Hochberg).

In contrast, abundant serotypes 16F and 19F in the nasopharynx were significantly prevalent among carriers, suggesting that they may have a lower potential to cause invasive disease. Other frequently detected serotypes, such as 6A, 6B, and 23F, were both abundant and evenly distributed among carriers and patients. It is conceivable that they might require a longer period of nasopharyngeal colonization compared to the hyper-invasive serotypes (1, 5, and 12F) before causing infections at sterile sites. Serotypes 6A, 6B, and 23F have previously been reported as common and abundant serotypes among both non-invasive carriers and those with invasive infections [38][39].

As mentioned earlier, the temporal distribution of the hyper-invasive serotypes (1, 5, and 12F) concerning the vaccine rollout timeline was noteworthy. The relative frequency of serotype 1 exhibited a significant increase (pre-PCV13=8.6% and post-PCV13=11.3%), while serotype 5 displayed a significant decrease (pre-PCV13=11% and post-PCV13=1.5%) following the introduction of the vaccination program in Malawi. This suggests that vaccination may be effective against serotype 5 but did not alleviate the burden of invasive pneumococcal diseases (IPDs) caused by serotype 1. Additionally, serotype 12F, which is not included in PCV13, showed a significant increase (pre-PCV13=1.1% and post-PCV13=5.7%) in the post-PCV13 era, indicating its potential emergence as an invasive strain. Nevertheless, a more extensive dataset, containing more recent samples, is essential to comprehensively characterize the long-term effects of PCV13.

### High diversity in the pneumococcal pan-genome

The genome assembly produced an average optimized assembly hash value of 96 and an average N50 of 113,986. The mean assembled genome size was estimated to be 2,116,779 nucleotides, with a standard deviation of 106,481. This length aligns within the range of previously reported *S. pneumoniae* genome sizes [40].

The pan-genome spanned 5,178,167 base pairs and encompassed 6,803 genes, comprising 729 core genes (10.7%), 820 soft-core genes (12.1%), and 5,254 accessory genes (77.2%). The pan-genome remained open, demonstrating a continuous increase in the number of genes as the sample size expanded (S5 Fig). The gene presence-absence heatmap in Fig 2 illustrates the pan-genome, revealing serotypes as the primary factor influencing gene distribution. Notably, distinct clustering was observed for hyper-invasive serotypes 1, 5, and 12F, forming unique clades. Specific sets of genes present in different serotypes were represented as distinctive blue blocks within the heatmap.

**Fig 2.**
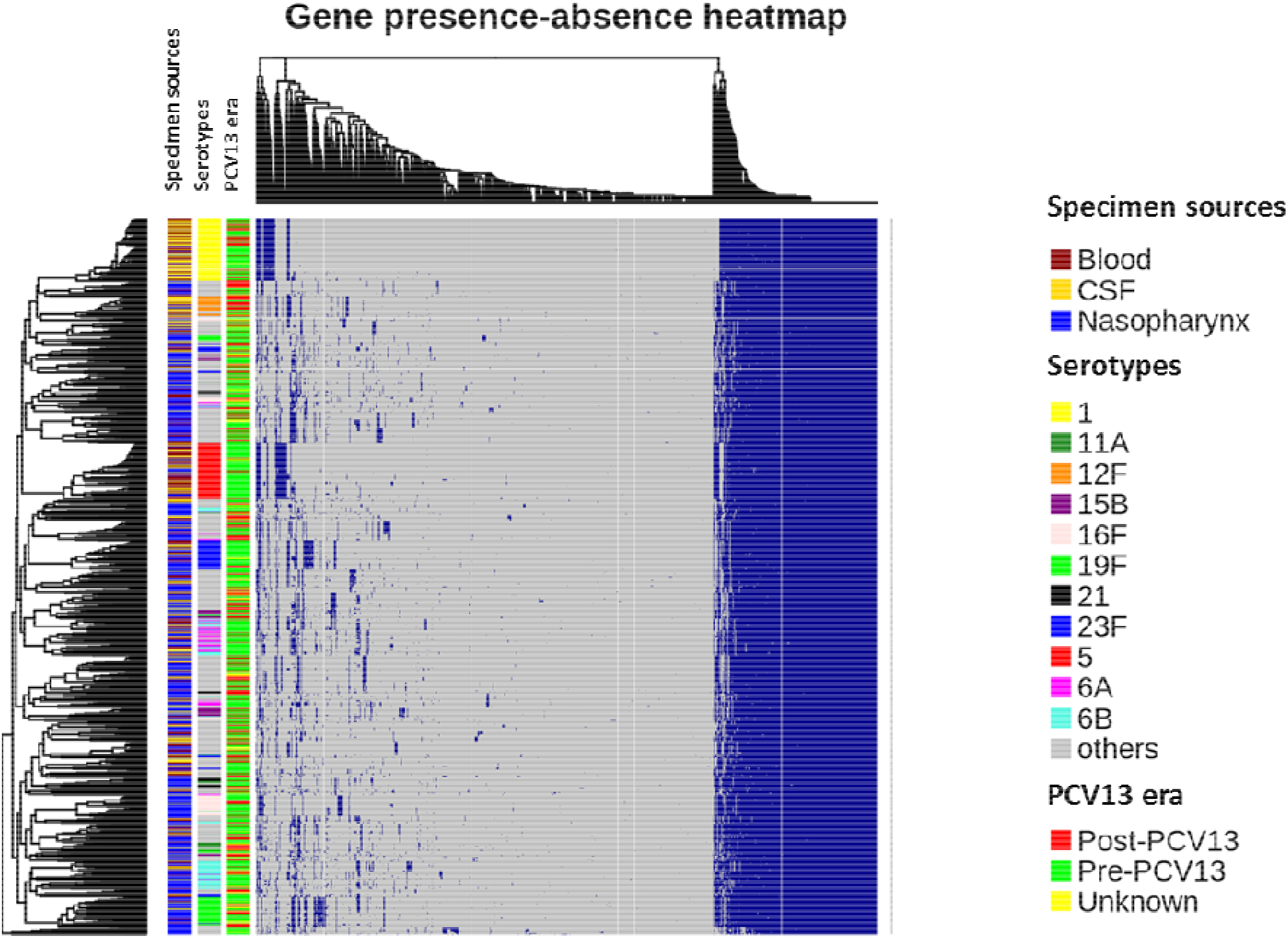
The pan-genome matrix of 1477 pneumococcal isolates from Malawi. The pan-genome is visualized as a gene presence-absence heatmap representing the hierarchical unsupervised clustering of samples based on the distribution of genes in the pan-genome. Each row is a sample, and each column is a gene. A blue dot denotes the presence of each gene. On the right side of the heatmap, the large blue block shows core genes present in all samples. The left side of the heatmap represents the accessory genome along with the clustering bands. In addition to the significant serotypes 1, 5, 12F, 16F, and 19F, other abundant serotypes, including 6A, 6B, and 23F, as well as serotypes with source-based p-value < 0.05, including 21, 11A, and 15B, are also highlighted on the heatmap.

### The significant invasive serotypes (1, 5, and 12F) showed the highest distinction in the core- and accessory-genome

The maximum-likelihood tree, depicting the distribution of small-scale variants (SNPs and indels) within the core-genome, highlighted the hyper-invasive serotypes 1, 5, and 12F as monophyletic clusters (Fig 3). These serotypes (1, 5, and 12F) were distinct, forming individual clusters that were more prominently separated compared to other abundant serotypes, such as 6B, 19F, and 23F, which appeared as multiple clusters on the phylogenetic tree (see Fig 3)

**Fig 3.**
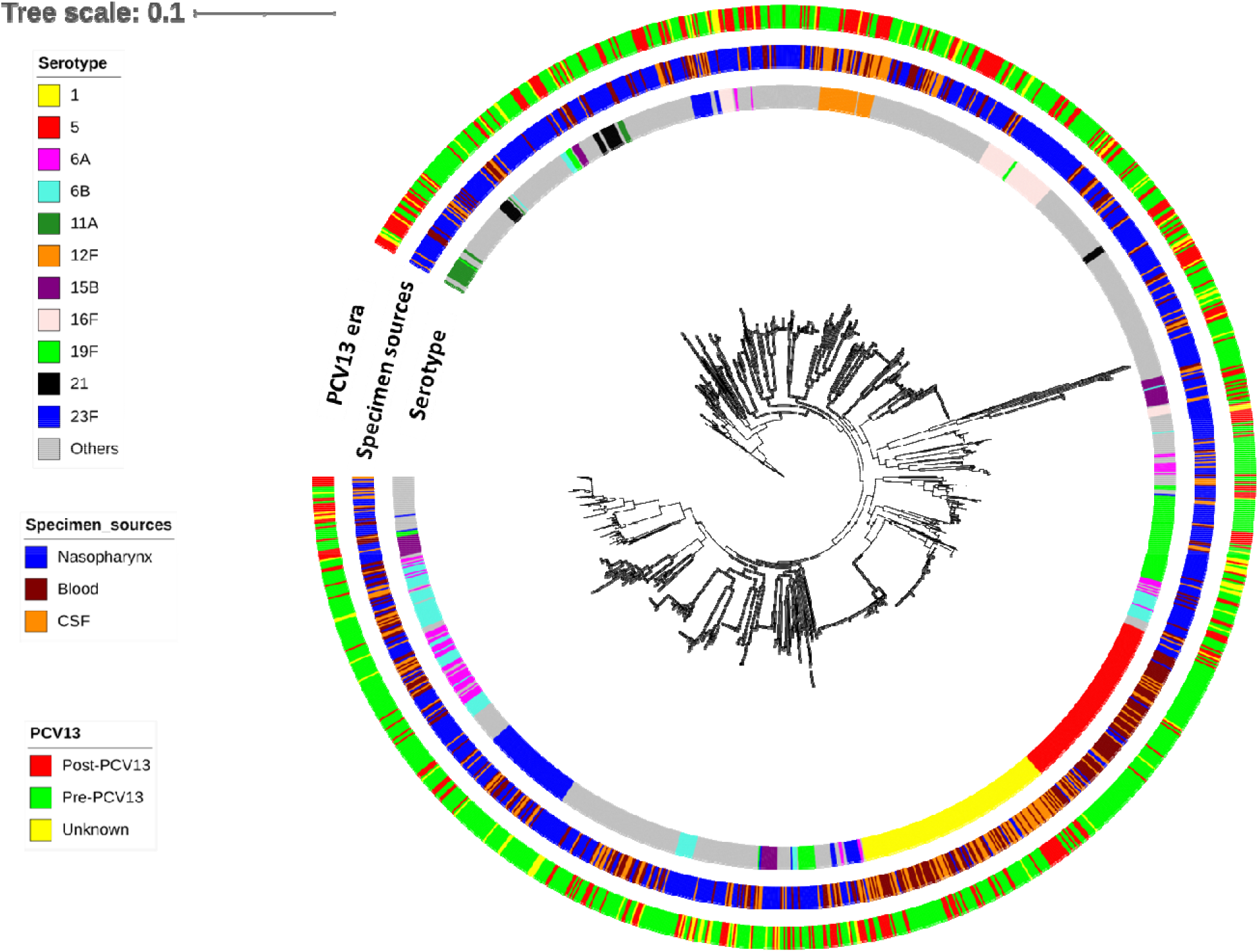
The phylogenetic population structure of 1477 pneumococcal samples from Malawi. The phylogenetic tree was built based on the multiple sequence alignment of the core genome using the maximum likelihood method. Colors on the loops show the serotypes, specimen sources (isolation sites), and PCV13 eras. In addition to the significant serotypes 1, 5, 12F, 16F, and 19F, other abundant serotypes, including 6A, 6B, and 23F, as well as serotypes with source-based p-value < 0.05, including 21, 11A, and 15B, are also highlighted on the tree.

The PCA of the large-scale variants in the accessory-genome (gene presence/absence) displayed serotypes 1 and 5 as distantly clustered from other strains (Fig 4). Additionally, a moderate level of separation was evident for serotypes 12F, 19F, and 23F. This distinct clustering of serotypes 1 and 5 is aligned with profiles demonstrated by the phylogenetic tree.

**Fig 4.**
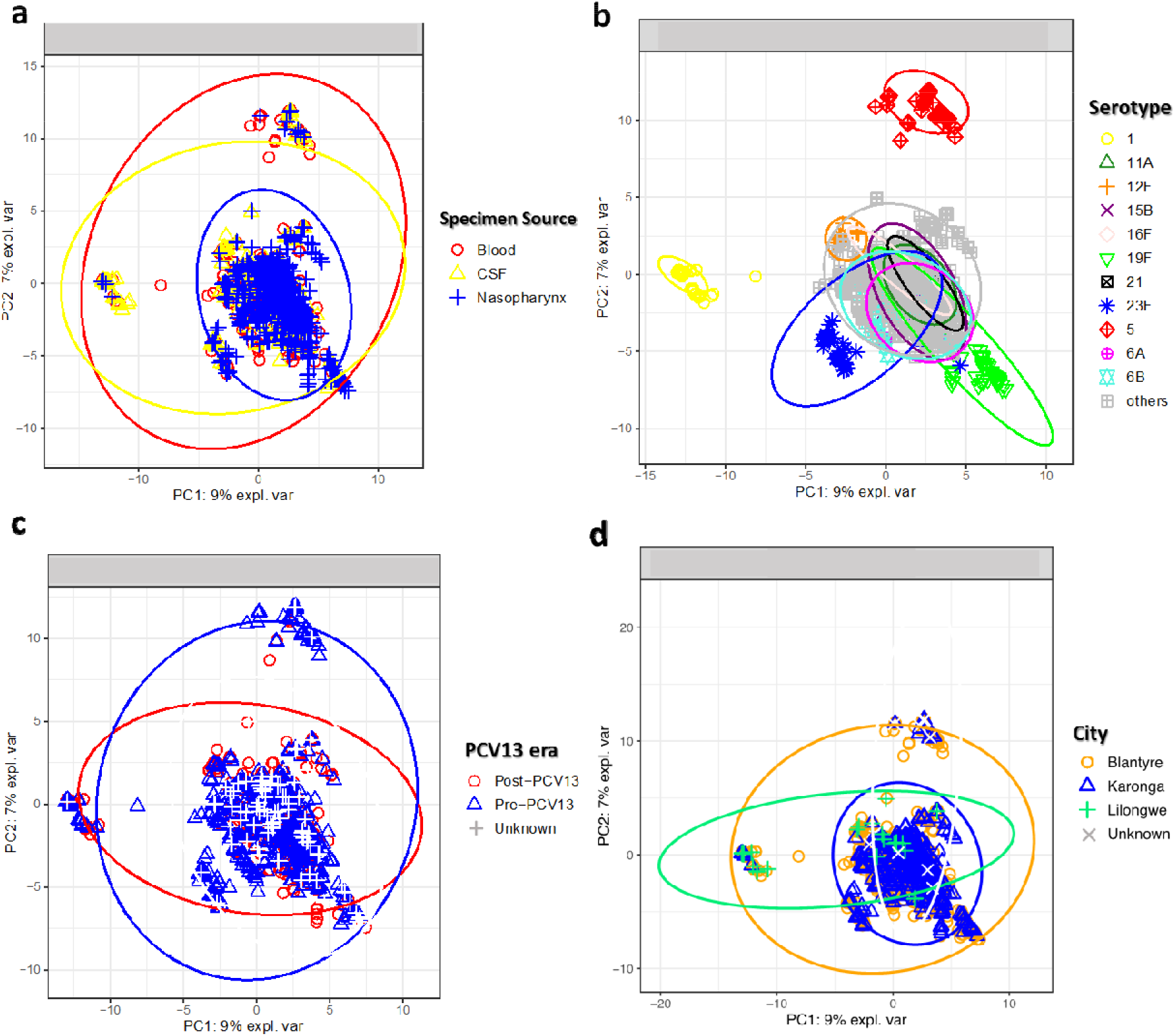
The PCA of the gene distribution in the pan-genome of pneumococcal isolates from 1477 Malawians. The PCA of variants (gene presence-absence) in the accessory-genome indicates the influence of (a) specimen sources (isolation sites), (b) serotypes, (c) PCV13 (vaccination) era, and (d) geographical locations on the gene presence-absence profile of pneumococcal isolates in Malawi. Serotypes 1 and 5 were clearly separated from other samples.

Figures 3 and 4 indicate that factors such as time, locations, and isolation sites (specimen sources) failed to sufficiently explain the small- and large-scale genetic variants in the pan-genome. Instead, the serotype of the samples emerged as the primary driver of the population structure. The hyper-invasive serotypes (1, 5, and 12F) exhibited the most significant core and accessory distances from other strains, signaling their genetic distinctiveness among disease-associated serotypes. This heightened genetic dissimilarity might be linked to their invasive potential.

It’s crucial to thoroughly consider population structure and assess any associations with disease across the population. An important observation is the absence of the same level of genetic distinction in serotype 12F, potentially due to its smaller sample count (n=35) compared to serotypes 1 (n=129) and 5 (n=116). Additionally, the near-complete separation of serotypes 1 and 5 from the others may primarily reflect their infrequent presence in the carriage group, suggesting that these patterns might stem more from sampling biases than genuine genetic variations. To address these concerns, ten samples from each hyper-invasive serotype and the PCV13 vaccine types were randomly selected from the nasopharynx, blood, and CSF. A PCA of the gene distribution was performed on the downsampled dataset, reiterated a noticeable pattern of clustering evident for serotypes 1, 5, and 12F each positioned far from other strains (S6 Fig).

Another aspect to consider revolves around the differentiation of serotypes. Theoretically, serotypes are differentiated due to their distinct serotype-defining capsule genes. However, a question emerges regarding the notable distinction in clustering observed among hyper-invasive serotypes (1, 5, and 12F) compared to other serotypes. Our investigation focused on the hypothesis that serotypes 1, 5, and 12F might have undergone gene acquisitions or losses, potentially contributing to their invasiveness. While there are other prevalent serotypes in blood and CSF, such as 6B and 23F, their similar prevalence in the nasopharynx suggests they might persist in the nasopharynx for extended durations compared to the hyper-invasive serotypes (1, 5, and 12F).

### The gene presence-absence statistical analysis

The following issues could skew the gene presence-absence analysis:

a. The batch effect introduced by geographical locations: 85% of carriage samples were from Karonga, whereas 95% of disease samples were from Blantyre (Table 1). A comparison between carriage and disease groups may only identify the difference between pneumococcal genomes from two geographical locations rather than between the non-invasive and invasive groups.
b. Study limitation: The likely presence of invasive serotypes in the carriage group made it unclear which nasopharyngeal samples progressed to disease after collection. Indeed, the carriage population likely contained invasive serotypes that could bias the test between carriage and disease samples to identify potential virulence genes.
c. Population structure: The significant abundance of the hyper-invasive serotypes (1, 5, and 12F) in the patient group and their highest genetic distinction would skew the test between the carriage and patient groups. The difference between carriage and disease groups would actually be the difference between the carriage and hyper-invasive serotypes (and not all strains in the blood and CSF).

To assess the geographical batch effect, carriage isolates from Karonga were compared with those from Blantyre. The gene presence-absence statistical test did not identify any significant genes differing between the Blantyre and Karonga groups, indicating a similarity in gene content within the carriage samples from both locations. Additionally, as previously observed, the serotype distributions in the carriage groups from Karonga and Blantyre displayed similarities (S3 Fig). Consequently, the impact of geographical location on the pneumococcal genomes was not substantial. This outcome aligns with expectations considering that Karonga and Blantyre are approximately 830 kilometers apart, and the demographic similarities between the populations in these cities.

While serotypes like 6A, 6B, and 23F, potentially associated with invasive traits, were found in both carriers and patients, comparing the entirety of the carriage and patient groups remains important. This is because serotypes identified as potential invasive in the nasopharynx might undergo genomic alterations before reaching sterile sites. During the colonization phase in the nasopharynx, these serotypes likely engage in genetic exchange via recombination and horizontal gene transfer with other pneumococci or bacterial species. Therefore, the genomic profile of an invasive serotype in the nasopharynx might differ from that in the blood and CSF.

To account for these complexities, our analysis involved an association test between the entire carriage and disease groups, excluding the hyper-invasive serotypes 1, 5, and 12F (a location-based analysis). This exclusion aimed to prevent these hyper-invasive serotypes from introducing biases when comparing the carriage and patient groups. The location-based analysis identified 27 significant genes, including 11 genes significantly present in the blood and CSF and 16 genes in the nasopharynx (Table 2, Table 3, and S2 Table for further details)

**Table 2.** Significant genes (p-value < 0.05) present in pneumococci in the blood and CSF (hyper-invasive serotypes 1, 5, and 12F were excluded) compared to the nasopharyngeal pneumococci.

| ID | Annotation | P-value | Odds ratio |
| --- | --- | --- | --- |
| SP_0360 | Capsular polysaccharide biosynthesis protein (from RD3) | 8.17E-05 | 4.897059 |
| SP_0358 | Capsular polysaccharide biosynthesis protein (from RD3) | 8.17E-05 | 4.897059 |
| SP_0351 | Capsular polysaccharide biosynthesis protein (from RD3) | 8.17E-05 | 4.897059 |
| SP_0359 | Capsular polysaccharide biosynthesis protein (from RD3) | 8.17E-05 | 4.897059 |
| SP_0357 | Capsular polysaccharide biosynthesis protein | 0.000175 | 4.758421 |
| SP_1037 | Type II restriction endonuclease Bcgl | 0.001074 | 2.878307 |
| SP_1953 | Bacteriocin/lantibiotic secretion ABC transporter permease protein | 0.001864 | 4.631892 |
| SP_0535 | Putative immunity protein | 0.004674 | 1.988523 |
| SP_1056 | Relaxase/Mobilisation nuclease domain (From RD6) | 0.012604 | 2.932153 |
| SP_1656 | Hypothetical protein | 0.014050 | 1.963212 |
| SP_0347 | Capsular polysaccharide biosynthesis protein (from RD3) | 0.020560 | 1.808997 |

**Table 3.** Significant genes (p-value < 0.05) present in the nasopharyngeal pneumococci compared to pneumococci in the blood and CSF (hyper-invasive serotypes 1,5, and 12F were excluded).

| ID | Annotation | P-value | Odds ratio |
| --- | --- | --- | --- |
| SP_1770 | Glycosyl transferase, glyB (from RD10) | 0.000132 | 0.502513 |
| SP_1771 | Glycosyl transferase, family 2/family 8 (from RD10) | 0.012770 | 0.560236 |
| SP_1763 | Preprotein translocase secY family protein (from RD10) | 0.017254 | 0.563256 |
| SP_1765 | Glycosyl transferase, glyF (from RD10) | 0.018150 | 0.564542 |
| SP_0939 | Hypothetical protein | 0.020740 | 0.483266 |
| SP_1766 | Glycosyl transferase, glyE (from RD10) | 0.020740 | 0.483266 |
| SP_1767 | Glycosyl transferase, glyD (from RD10) | 0.023196 | 0.568879 |
| SP_1762 | Accessory secretory protein asp1 (from RD10) | 0.023196 | 0.568879 |
| SP_1761 | Accessory secretory protein asp2 (from RD10) | 0.023196 | 0.568879 |
| SP_1760 | Accessory secretory protein asp3 (from RD10) | 0.023196 | 0.568879 |
| SP_1755 | Hypothetical protein | 0.031054 | 0.558554 |
| SP_1757 | Glycosyl transferase, glyB (from RD10) | 0.031054 | 0.558554 |
| SP_1764 | Glycosyl transferase, glyG (from RD10) | 0.023196 | 0.568879 |
| SP_1768 | Conserved hypothetical protein (from RD10) | 0.031054 | 0.558553 |
| SP_1758 | Poly(glycerol-phosphate) alpha-glucosyltransferase, tagE (from RD10) | 0.031122 | 0.571941 |
| SP_1759 | Preprotein translocase, secA subunit (from RD10) | 0.031321 | 0.574557 |

The most significant genes identified in both blood and CSF belonged to the cps locus (RD3), suggesting a potentially increased level of encapsulation during disease. Specifically, genes SP_0357, SP_0358, and SP_0360 encode epimerases involved in the biosynthesis of complex lipopolysaccharides, which are essential components of the pneumococcal capsule. SP_0351 encodes a membrane protein glycosyltransferase responsible for catalyzing glycosyl group transfer during capsule synthesis, and SP_0359 encodes UDP-2-acetamido-2,6-beta-L-arabino-hexul-4-ose reductase, a crucial protein involved in capsular polysaccharide biosynthesis. Other significant genes in the blood and CSF, including SP_1953, SP_0535, SP_1037, and SP_1056, are involved in toxic secretion and recombination. SP_1056 is part of the pneumococcal pathogenicity island 1 (PPI1) located within RD6. This gene encodes a mobilization protein necessary for the horizontal transfer of genes and plasmids via bacterial conjugation. SP_1056 plays a role in forming the relaxation complex or relaxosome by interacting with other enzymes [41].

Significant genes identified in the nasopharynx (absent in samples from blood and CSF) originated from RD10, recognized as the SecY2A2 island responsible for the secretion of pneumococcal serine-rich repeat protein (PsrP) (S7 Fig) [42]. RD10 contains several glycosyltransferases and secretory components that were significantly missing from the genome of samples obtained from blood and CSF.

To explore the divergence of the hyper-invasive serotypes (1, 5, and 12F), they were compared to serotypes 16F and 19F, which were significantly present in the nasopharynx (serotype-based analysis). Serotypes 16F and 19F could represent non-invasive strains better than the whole nasopharyngeal population that potentially contained some invasive serotypes. Indeed, it was a test between serotypes with the highest and lowest invasiveness to characterize the genomes of serotypes 1, 5, and 12F. The gene gain and loss profiles in the hyper-invasive serotypes may include components contributing to their virulence and short colonization period.

The serotype-based analysis identified 184, 157, and 186 significant genes (present/absent) in serotypes 1, 5, and 12F, respectively (S3 Table, S4 Table, and S5 Table) that were much larger than the number of significant genes identified by the location-based analysis. The functional enrichment analysis identified the phosphotransferase system (PTS, KEGG ID: spn02060) as over-represented and oxidative phosphorylation (KEGG ID: spn00190) as under-represented pathways in the hyper-invasive serotypes 1, 5, and 12F (p-value < 0.05).

In total, there were 18 significant genes jointly present in the hyper-invasive serotypes (Fig 5.a), including elements of the PTSs that transport sucrose and lactose across the membrane (SP_0302, SP_0303, SP_0304, SP_0305, SP_0306, SP_0308, SP_0309, and SP_0310), bacteriocins (SP_0544 and SP_1051) and a permease protein (SP_1527). The over-represented pathway (spn02060) was associated with significant genes that code for PTS transporters involved in carbohydrate metabolism. Seven genes were unannotated. The PTS transporters genes were also present in a high proportion of abundant serotypes in sterile sites such as 6B (67%) and 23F (86%).

**Fig 5.**
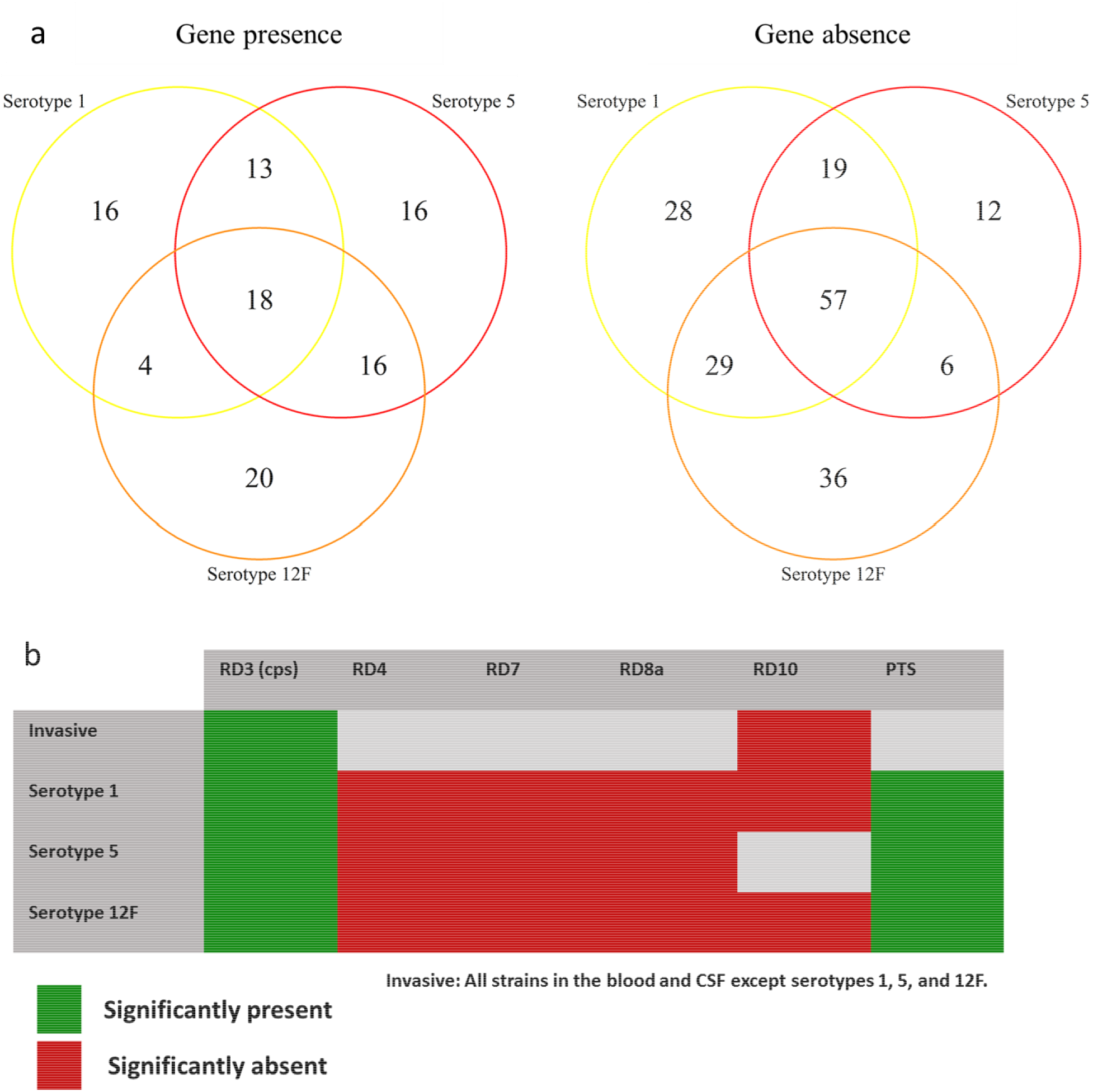
The summary of the gene presence-absence analysis. (a) The number of significant genes present-absent in the hyper-invasive serotypes. The gene presence-absence analysis was applied using Scoary to compare the gene pools of the hyper-invasive serotypes and serotypes 16F and 19F. P-values were corrected by the Bonferroni method, and significant genes had an adjusted p-value of less than 0.05. (b) The significant presence and absence of RDs in samples from blood and CSF is shown as a presence-absence heatmap.

A total of 57 significant genes were absent in serotypes 1, 5, and 12F (Fig 5.a). The most significant absences were observed within RD8a, consisting of two operons, RD8a1 (SP_1315-1324) and RD8a2 (SP_1325-SP_1331) (S8 Fig). RD8a1 harbors eight *ntp* genes, that code *V-type proton/sodium ATP synthase complex* that produces ATP via oxidative phosphorylation in the presence of a Na+ gradient across the membrane[43]. RD8a2 includes *neuraminidase*, *N-acetylneuraminate lyase (nanA),* and *N-acetylmannosamine-6-phosphate epimerase (nanE).* These genes cleave carbohydrates from the glycoproteins on the surface of epithelial cells. Other genes in RD8a2 encode the Sodium/solute symporter subunits that use Na^+^ gradient to import the carbohydrates [44]. Symporter refers to a channel that transports the solute (carbohydrates) and co-solute (Na+) in the same direction by utilizing the energy stored in an inwardly directed sodium gradient. Fundamentally, the genes within RD8a operons collaborate to generate ATP, cleave carbohydrates from the host epithelial cells, and import them into the bacterial cell. RD8a was absent in all samples associated with serotypes 1, 5, and 12F, while it was present in other prevalent serotypes like 6A, 6B, 16F, and 19F. The pathway sp00190, which was underrepresented in hyper-invasive serotypes, was associated with genes within RD8a. The absence of RD8a in hyper-invasive serotypes (1,5, and 12F) may be linked to their rapid invasion into the blood and CSF, where the availability of free oxygen molecules necessary for oxidative phosphorylation is limited [45].

Other significant genes absent from the hyper-invasive serotypes were from RD4 and RD7. RD4 consists of a cluster of sortase enzymes responsible for the assembly of pilins into pili and anchoring these structures and other surface proteins to the cell wall [46][47]. The pilus is a hair-like structure associated with bacterial adhesion and colonization [48]. Owing to the hypothesized short colonization period of hyper-invasive serotypes, they may not harness the benefits of RD4 genes involved in pilus assembly. RD7 genes remain uncharacterized as of the present date.

It is worth mentioning that there were similarities between hyper-invasive serotypes (1,5, and 12F) and other strains in the blood and CSF. As observed for samples in the blood and CSF (Table 2), capsule genes were also significantly present in the hyper-invasive serotypes (1,5, and 12F). Moreover, RD10 (previously found to be significantly absent from blood and CSF as described in Table 3) was also absent from serotypes 1 and 12F. However, RD10 was fully conserved in serotype 5. RD10 was also conserved in 100% of serotypes 16F and 19F. The summary of the gene present-absent analysis is illustrated in Fig 5.b.

Finally and as mentioned, serotypes 6B and 23F were abundant in both carrier and patient groups, the intra gene presence/absence statistical test for serotype 6B between the nasopharynx (n=60) and sterile sites (n=37), and for serotype 23F between the nasopharynx (n=43) and sterile sites (n=50) did not identify any significant gene. The gene content of these two serotypes in the nasopharynx and sterile sites was similar.

To highlight genes that might assist pneumococci in crossing the blood-brain barrier, a test was also conducted between the samples from blood and CSF. The analysis between whole blood (n=368) and CSF (n=284), serotype 1 samples from blood (n=60) and CSF (n=61), and serotype 5 samples from blood (n=75) and CSF (n=23) did not identify any significant genes.

## Discussion

The *S. pneumoniae* genome is highly diverse, with only a small portion of genes conserved across all strains. In this species, the pan-genome is open, allowing for an extensive gene repertoire due to the highly recombinogenic nature of pneumococci. Changes in the *S. pneumoniae* habitat may lead to the utilization of various gene combinations, enabling organisms to diversify their genome and respond effectively to environmental stresses. This study found a high genetic diversity, with merely 10.7% of genes classified as core. These core genes have been conserved across all samples for an extensive period, at least from 1997 to 2015. Their presence may be crucial for cell survival, making them potential targets for drug design and vaccine development. Specifically, core genes without any SNPs in their structure are of particular interest. Notable conserved core genes identified in this study included SP_1961 (rpoB, DNA-dependent RNA polymerase), SP_0251 (formate acetyltransferase), SP_1891 (amiA, Oligopeptide binding protein), and SP_0855 (parC, Topoisomerase IV). These conserved core genes play integral roles in DNA transcription and translation.

Serotypes 1, 5, and 12F exhibited a high prevalence among patients but were rarely found in the carrier group. This observation indicates their increased invasiveness, likely due to a short duration of nasopharyngeal colonization. Conversely, serotypes 16F and 19F were significantly more frequent among carriers, suggesting their dominance in the nasopharynx but with lower invasiveness. Most other serotypes were common among both carriers and patients. Knowing that pneumococcal virulence strongly depends on the serotype of isolates, we sought to address why several serotypes were shared across both nasopharynx and sterile sites. Here, we discuss two possible scenarios that could justify the ubiquitous presence of some serotypes in both nasopharynx and sterile sites.

The first scenario is related to the colonization of *S. pneumoniae*, which is known as a prerequisite for virulence[4]. Several samples from the carriage group may be actually the invasive serotypes collected during their colonization phase. Abundant serotypes such as 6B and 23F that had a similar frequency amongst carriers and patients need to colonize the upper respiratory tract longer than the hyper-invasive serotypes before entering the sterile organs. In contrast, the hyper-invasive serotypes 1, 5, and 12F colonize the nasopharynx for a short period and quickly enter the sterile sites.

The second possible scenario relates to the differential gene expression pattern of shared genes in the ubiquitous serotypes. Although the type-specific *cps* genes were identified in the isolates of both nasopharynx and sterile sites, the expression pattern of these genes could vary within each strain, which would contribute to the invasiveness of some strains. Several studies described a cycle of encapsulation and un-encapsulation amongst pneumococcal strains. Isolates benefit from mutations in the *cps* locus to either cease or re-start the capsule expression [49]. The lack of a capsule at the epithelial surface enables the bacterium to expose its surface proteins on the cell wall underneath the capsule and promote adherence to the host epithelial cells. It has been estimated that 15% of isolates in the upper respiratory tract are unencapsulated and adhere to the respiratory epithelial cells more efficiently than encapsulated isolates [50][51]. Lack of capsule also facilitates acquiring virulence and resistance genes from other isolates. The thick capsule prevents immunoglobulins from interacting with the pathogen surface proteins during disease. Meanwhile, the negatively charged CPS interferes with the function of the host phagocytes [52][53]. Taken together, the presence of the *cps* locus in the genome of isolates assigned to the same serotype does not necessarily reflect the encapsulation of all cells.

Serotypes 1 and 5 can infect all age groups and cause severe IPDs [54]. Serotype 1 is genetically distinct between different geographical regions [55] and is known as the leading cause of pneumococcal meningitis in Africa [56][57]. Our findings support previous research showing that serotypes 1, 5, and 12F are the major cause of IPDs in Malawi [58]. Serotype 1 was persistently dominant in pre- and post-PCV13 eras, serotype 5 was only predominant in pre-PCV3, and serotype 12F emerged after vaccination. The study also characterized the high genetic distinction of the hyper-invasive serotypes in Malawi by identifying significant genes that are present or absent in their genome structure compared to other serotypes. Many of the significant genes present in the nasopharynx and sterile sites were homologous and had the same function, and there were significant genes with an unknown function (hypothetical proteins). However, of greatest interest, we did find several significant genes with specific functions that could explain the difference between the biology of the hyper-invasive serotypes and nasopharyngeal samples.

Genes within RD8a were significantly absent in the genome of serotypes 1, 5, and 12F. RD8a is known as a region previously linked to the virulence of serotypes 6B and 14 in the United States [59]. Our study identified this region’s conservation in serotypes 13, 14, 16F, and 19F in Malawi. This observation prompts the hypothesis that RD8a may be essential for the prolonged colonization of serotypes contrasting with the quicker colonization of serotypes 1, 5, and 12F. The functions of genes in RD8a strengthen the assumption to some extent that RD8a may be essential for long nasopharyngeal colonization. The genes within RD8a, such as neuraminidase, *nanA*, and *nanE*, are involved in the cleaving of terminal sialic acid residues from mucoglycans and epithelial glycoconjugates. This activity aids the pathogen in breaching the mucus layer and adhering to the epithelial cells in the nasopharynx. Additionally, since free carbohydrates are limited in the upper respiratory tract [60], cleaved sialic acid can be used as the carbon source for metabolism. Moreover, during colonization, secretion of the pneumococcal toxins elevates the level of sodium ions (Na^+^) in the nasopharynx [61], which enables the sodium-solute symporter in RD8a to import a wide variety of substrates with the sodium ions into the cell [62]. Most importantly, the *ntp* gene cluster in RD8a encodes the *V-type sodium ATP synthase* that pumps the extra sodium ions out of the cell [43] and uses the sodium-motive force for oxidative phosphorylation and ATP synthesis [63]. Oxidative phosphorylation is the final step of aerobic respiration that requires free oxygen molecules for ATP synthesis. Pneumococci are facultative anaerobes that can either perform aerobic or anaerobic respiration with or without oxygen. In the upper respiratory tract, they access atmospheric oxygen molecules that can be used by *ntp* genes to perform oxidative phosphorylation. However, genes in RD8a may not be beneficial for hyper-invasive serotypes that supposedly do not stay in the nasopharynx for long. Thus, the level of aerobic ATP synthesis is presumably higher in serotypes 13, 14, 16F, and 19F in contrast with serotypes 1, 5, and 12F, which lack RD8a.

Pneumococci can ferment up to 30 types of carbohydrates, imported mainly by two types of membrane transporters, including ATP-binding cassette (ABC) transporters and PTS transporters [64]. The major differences between ABC and PTS transporters are: (i) ABC transporters use energy from ATP, but PTS transporters use energy from phosphoenolpyruvate, and (ii) ABC transporters do not modify the imported substrate, but PTS transporters phosphorylate the incoming sugar upon transport. Generally, ABC transporters require more energy than PTS transporters, albeit they can transport longer and more complicated carbohydrates [65]. Unlike isolates in the nasopharynx, serotypes 1, 5, and 12F have access to more simple and free host dietary carbohydrates in the blood and the central nervous system. Due to a potential lower ATP synthesis level in serotypes 1, 5, and 12F (due to the lack of RD8a), they may prefer to use PTS transporters to uptake sugars such as fructose and lactose, and that is why genes that encode PTS transporters are significantly more present in the hyper-invasive serotypes (1, 5, and 12F). In addition to the sugar uptake, PTS transporters regulate several pathways in bacteria, such as gene expression and communication between cells. Thus, the phenotypic effects of the PTS transporters should not be limited just to their ability to import carbohydrates [66].

Genes within RD10 were absent from serotypes 1 and 12F. However, they were conserved in serotypes 5, 16F, and 19F. Moreover, the location-based gene presence-absence analysis showed that RD10 was significantly present in nasopharyngeal samples in comparison to samples collected from sterile sites. Operon RD10 in pneumococcus shares homology with the general secretion pathway protein B sceA2/Y2 system components in Streptococcus gordonii, which are involved in secreting the general secretion pathway protein B linked to infective endocarditis [67]. In the*S. pneumoniae* genome, the homolog of general secretion pathway protein B is PsrP, which is transported to the bacterial cell surface by the SecA2/Y2 system encoded by genes in RD10. Research on *Streptococcus gordonii* has indicated that the presence of SecA2/Y2 facilitates adhesion to both epithelial cells in the nasopharynx and erythrocytes in the blood. [68][69]. This may explain why SecA/Y2 is significantly present in nasopharyngeal samples and serotype 5 (abundant in the blood). The presence of the secA2/Y2-like component should also facilitate the export of pneumolysin, which enhances adhesion to the host cell and contribute to survival in the blood [70][71].

In conclusion, specific genes present or absent in the hyper-invasive serotypes (1, 5, and 12F) may play a role in their invasiveness and lower colonization rate. Nonetheless, experimental validation is necessary to confirm the computational findings from this study. While the serotype is the primary determinant of the pneumococcal population structure, this research has highlighted the substantial genetic divergence of serotypes 1, 5, and 12F compared to other serotypes. Their substantial presence in the blood and CSF accounted for the most pronounced genomic and functional differences observed between the nasopharynx and sterile sites. The lower frequency of serotypes 1, 5, and 12F among carriers could be attributed to their shorter colonization duration before entering sterile sites. These invasive serotypes possess elements of PTS transporters but lack genes from RD8a. Interestingly, RD10 is highly conserved in serotype 5, while it is absent in serotypes 1 and 12F. Notably, this study demonstrates that isolation sites do not significantly influence the genomic structure of pneumococcal isolates. Although a few genes were linked to the virulence of commonly present serotypes in both the nasopharynx and sterile sites, it is suggested that other high-throughput techniques like gene expression analysis may reveal the differences between these isolates more comprehensively. In summary, this research sheds light on the pneumococcal population structure and serotypes in Malawi. The unique cluster of significant genes in the hyper-invasive serotypes, along with highly conserved core genes, could serve as potential therapeutic targets.

## Author contributions

Sample collection, metadata curation, and genome sequencing: Jennifer Cornick, Dean Everett, Anmol Kiran, Chrispin Chaguza, and Chikondi Peno.

Methodology and data analysis: Arash Iranzadeh, Anmol Kiran, and Arghavan Alisoltani.

Result interpretation: Arash Iranzadeh, Arghavan Alisoltani, Nicola Mulder, and Dean Everett. Initial manuscript writing: Arash Iranzadeh.

Review of the manuscript: Arash Iranzadeh, Arghavan Alisoltani, Anmol Kiran, Robert F Breiman, Chrispin Chaguza, Chikondi Peno, Dean B Everett, Nicola Mulder.

All authors have given consent to participate in the study.

## Acknowledgment

Computations were performed using facilities provided by the University of Cape Town’s ICTS High-Performance Computing team: hpc.uct.ac.za. The authors also acknowledge the Centre for High-Performance Computing (CHPC), South Africa, for providing computational resources to this research project. We thank the study participants and all involved staff at the Karonga Prevention Study and the Malawi-Liverpool-Wellcome Trust Clinical Research Programme. We thank Olivier Koole and Naor Bar-Zeev for their scientific input.

## Ethical approval

Not required as the research delas with bacterial samples.

## Supporting information

**S1 Fig. Characteristics of the 1477 pneumococcal isolates used in the study.** (a) The relative frequency of serotypes in the entire cohort, samples were assigned to 56 serotypes. For each sample, the in-silico serotyping was accomplished by SeroBA. (b) Frequency of isolates in the pre- and post-PCV13 eras in Malawi. (c) Frequency of isolates obtained from each specimen source.

**S2 Fig. Distribution of the abundant serotypes (frequency > 5%) before and after the vaccination rollout in Malawi in 2011**. Serotype 1 persistently dominated both the pre- and post-vaccination eras.

**S3 Fig. The serotype distribution among carriers in Karonga and Blantyre.** Distributions were similar, except for serotype 6B, which was more dominant in Karonga, and serotype 13, which was more prevalent in Blantyre.

**S4 Fig. Serotype distribution among meningitis patients in Lilongwe and Blantyre.** Only 3.5% of disease samples (23 out of 652, i.e., 3.5%) were collected from Lilongwe. Serotypes 1 and 12F were predominant in both regions; however, a larger dataset from Lilongwe is needed to accurately reflect the true serotype distribution in this area.

**S5 Fig. The pan-genome of 1477 pneumococcal samples isolated in Malawi was obtained from 1997 to 2015.** The pan-genome is an open pan-genome, which means the number of total genes increases unlimitedly when the sample size grows. The dashed line represents the number of total genes, and the solid line represents the number of conserved genes in the pan-genome.

**S6 Fig. The three-dimensional PCA of the gene distribution in the vaccine types.** For each serotype and for downsampling, 10 samples were randomly selected from the nasopharynx, blood, and CSF. The PCA was conducted using the R package MixOmics. Hyper-invasive serotypes 1, 5, and 12F clustered separately from other strains.

**S7 Fig. Genes in RD10 are absent from serotypes 1 and 12F but conserved in serotype 5, 16F, and 19F.** Genes from RD10 encode the components of the secretory system SecA2/Y2 that transports glycoproteins to the bacterial cell surface, which are required for binding to the human proteins on the surface of epithelial cells and erythrocytes.

**S8 Fig. RD8a consists of two operons RD8a1 (SP_1315-1324) and RD8a2 (SP_1325-1331).** This region is not detected in the significant invasive serotypes 1, 5, and 12F, but it is present in more than 80% of serotype 16F and 19F that significantly dominates the nasopharynx. The important biological processes carried out by these genes are the transport of ions across the membrane and the synthesis of ATP molecules.

**S1 Table. Statistical analysis of serotypes’ prevalence across specimen sources.**

**S2 Table. Gene presence-absence analysis (Invasive vs Nasopharyngeal, serotypes 1, 5, and 12F were excluded).**

**S3 Table. Gene presence-absence analysis (Serotype 1 vs 16F & 19F). S4 Table. Gene presence-absence analysis (Serotype 5 vs 16F & 19F). S5 Table. Gene presence-absence analysis (Serotype 12F vs 16F & 19F).**

**S4 Table. Gene presence-absence analysis (Serotype 5 vs 16F & 19F).**

**S5 Table. Gene presence-absence analysis (Serotype 12F vs 16F & 19F).**

## Data summary

**S6 Table. Samples IDs on European Nucleotide Archive (ENA)**

